# Pooled enrichment sequencing identifies diversity and evolutionary pressures at NLR resistance genes within a wild tomato population

**DOI:** 10.1101/040998

**Authors:** Remco Stam, Daniela Scheikl, Aurélien Tellier

**Author notes:** Author for Correspondence: Remco Stam [Data Deposition: Data for this project has been deposited under NCBI BioSample SAMN04506098].

## Abstract

Nod-like Receptors (NLRs) are Nucleotide-binding domain and Leucine rich Repeats (NB-LRR)-containing proteins that are important in plant resistance signaling. Many of the known pathogen Resistance (R)-genes in plants are NLRs and they can recognise directly or indirectly pathogen molecules. As such, divergence and copy number variants at these genes is found to be high between species. Within populations, positive and balancing selection are to be expected if plants coevolve with their pathogens. In order to understand the complexity of R-gene coevolution in wild non-model species, it is necessary to identify the full range of NLRs and infer their evolutionary history.

Here we investigate and reveal polymorphism occurring at 220 NLR genes within one population of the partially selfing wild tomato species *S. pennellii*. We use a combination of enrichment sequencing and pooling of ten individuals, to specifically sequence NLR genes in a resource and cost-effective manner. We focus on the effects which different of mapping and SNP calling software and settings have on calling polymorphisms in customized pooled samples. Our results are accurately verified using Sanger sequencing of polymorphic gene fragments. Our results indicate that some NLRs, namely 13 out of 220, have maintained polymorphism within our *S. pennellii* population. These genes show a wide range of π_*N*_/π_*s*_ ratios and differing site frequency spectra. We compare our observed rate of heterozygosity to expectations for this selfing and bottlenecked population. We conclude that our method enables us to pinpoint NLR genes which have experienced natural selection in their habitat.

## Introduction

Resistance genes are important players in the interaction between plants and pathogens. They are involved in direct and indirect recognition of effector molecules from the pathogen and are hence thought to be under constant evolutionary pressure.

Most resistance genes (hereafter R-genes) including the best characterised ones belong to the NLR (Nod-like receptors) or NB-LRR (Nucleotide Binding site and Leucine Rich Repeat containing) type (Caplan et al. 2008). These include important R-genes from many food crops like Bs2 in pepper (Tai et al. 1999), R3a in potato (Huang et al. 2004) and Mi in tomato (Rossi et al. 1998). All NLR genes code for receptor proteins with a Nucleotide Binding Site (NB) and C-terminal Leucine Rich Repeats (LRR). Generally these NB-LRRs can be divided into two groups, based on the sequence of their NB-ARC domain and their N-terminal domains. One group has N-terminal domains related to the Toll and Interleukin Receptors (TIR), whereas the second non-TIR group often contains a Coiled Coil (CC) domain (McHale et al. 2006).

Resistance conferred by R-genes was thought to predominantly come from direct gene-for-gene interaction between the R-gene and pathogen avirulence effectors (avr) (Flor 1971). This recognition results in a strong defence response, called effector triggered immunity (ETI), which in place results in the production of reactive oxygen species or a hypersensitive response in the plant. This reaction leads to localised cell death and thus stops the spread of the pathogen (Morel & Dangl 1997). Several indirect modes of action have also been described. In these cases, NLRs detect the modification of a (guarded) target protein which triggers a similar defence response (Van der Biezen & Jones 1998; McHale et al. 2006). Several examples exist that confirm direct interactions (e.g Dodds et al. 2006), even though few known sites for direct interaction are known.

R-gene effector interaction might also be more complex. In wheat Lr10 and RGA, both NLRs, need to be present simultaneously to confer leaf rust resistance (Loutre et al. 2009). In tomato NRC proteins are required for resistance conferred by several other NLR (Gabriels et al. 2007; Wu et al. 2015). When over-expressed in planta, individual domains of Rx, a tobacco virus NLR, interact with each other (Moffett et al. 2002) rice multiple NLRs and their various combinations have been linked to highly redundant resistance profiles (Zhang et al. 2015).

The effector-NLR interactions are crucial to determine the outcome of infection. NLRs are therefore expected to show variations and evidence of selective pressures. In this light, NLRs are often found as large gene families and consequently annotation, origin and evolution of NLRs in plants (and animals) is an important field of study (Jacob et al. 2013; Maekawa et al. 2011). The numbers of identified NLR differ greatly within and between plant families, but also based on annotation methods. In the *Arabidopsis thaliana* about 150 NLR genes have been identified (Meyers et al. 2003). In Solaneceous species like tomato and potato this number rises to about 355 and 438 respectively (Jupe et al. 2012; Andolfo et al. 2014). In rice so far 466 NLRs have been annotated (Li et al. 2010). No clear correlations seem to exist between age, genome size and number of NLR since for example in the brassica family *Brassica rapa* which has a similar sized genome to *A. thaliana*, has only 80 known NLRs (Mun et al. 2009).

In *Arabidopsis thaliana*, NLR genes are located clusterwise on the genome and due to their hypervariable nature a model of a rapid birth and death process was suggested to explain expansion and diversification of the gene family (Michelmore & Meyers 1998). The 150 NLRs identified in A*rabidopsis thaliana* are very divergent, but it is possible to cluster many of them together in groups by sequence similarity, while some remain orphan. Of the 22 groups, 10 groups show genes with positively selected positions. The number of sites however varies from 1 to 26 and whereas the majority of selected sites occur in the LRR region, still 33 out of 116 are located in the NBS domain or other regions (Mondragón-Palomino et al. 2002). Studies of worldwide within-species variability of NLRs demonstrated the strong pervasive selection pressure. NLRs are thus likely to evolve under neutrality or purifying selection, and few under balancing selection (Bakker et al. 2006; Stahl et al. 1999). A study including sequence data from both *A. thaliana* and *A. lyrata*, showed similar results using divergence estimates, and indicated that the genes unique to a species, e.g. lacking homologues, appeared to show weaker selective pressure and less copy number variation (Guo et al. 2011).

Other studies focused on comparing the NLR complement between multiple species, and 2,363 NLRs were identified in 12 eudicot plants, including six crop species. Of these genes, 50% show tandem duplications associated with strong positive selection (the ratio of non-synonymous to synonymous substitutions, Ka/Ks > 1.5). However, a small set of NLRs appears to be conserved for over 100 MY in most eudicot genomes (Hofberger et al. 2014). In monocots the divergence between species appears to be large, as numbers of NLRs differ greatly between maize, sorghum, brachypodium and rice (Li et al. 2010). NLR clusters build from phylogenetic methods can exhibit a wide range of Ka/Ks ratios (0.5 – 3.3) (Yang et al. 2013). Since between species comparisons have lower statistical power to detect selection if divergence is high (Gharib & Robinson-Rechavi 2013), and they do not allow detecting the occurrence of balancing selection, we investigate within population variation to understand short-term evolution of NLRs.

Wild Solanum species provide the optimal model organisms for such studies. During its domestication *S. lycopersicum* has suffered significant from a reduction in genetic diversity (The 100 Tomato Genome Sequencing Consortium et al. 2014; Lin et al. 2014). Hence, wild tomato species regularly serve as germplasm source in current breeding programmes, making them economically interesting to study (Bai & Lindhout 2007). In addition, genomic resources are already available for a selection of wild and cultivated tomato.

In this study we make use of *S. pennellii*. This wild species contains several disease resistance loci, including canonical NLRs, against Oomycete pathogen *P infestans*, (Smart et al. 2007). It is the source for the I-1 and I-3 genes which confer resistance against Fusarium wilt (Sarfatti et al. 1991; Scott et al. 2004). It also contains other resistance loci, like RXopJ4, a bacterial spot Resistance locus (Sharlach et al. 2012) and has thus large value for plant breeders. *S pennellii*, LA0716 has been used to develop introgression lines with *S. lycopersicum* cultivar M82, which have been instrumental in understanding yield parameters and generating increased yields (Eshed & Zamir 1994; Eshed et al. 1996; Gur & Zamir 2004). *S. pennelli* is a self-compatible species which is expected to show low levels of within-population diversity. The recent sequencing of one plant of *S. pennellii* LA0716 yielded a high quality reference genome and led to the identification a number of abiotic stress associated genes (Bolger, Scossa, et al. 2014).

The costs of generating NGS data is constantly dropping, however, for complex plant species with large genomes, sequencing costs and also computation time for mapping or assembly are still considerable. R-gene enrichment sequencing can be used to reduce the complexity of the DNA sample, by enriching the R-gene component and thus reducing overall sequence complexity before sample submission. To this purpose RENSeq has successfully used to identify the NLR complement of both cultivated tomato and potato (Andolfo et al. 2014; Jupe et al. 2013). Nevertheless, for population genetics studies, ideally large numbers of individuals per population as well as large numbers of populations are desired to allow inference of short time scale selective pressures, and thus driving up in return the sequencing costs. Recently several studies have shown that pooled sequencing can dramatically reduce the sequencing costs, as well as time and costs associated with sample preparation (Schlötterer et al. 2014). Note that with pooled sequencing it is not possible to assign sequences to a single individual, but population genetics statistics can be successfully computed (Ferretti et al. 2013) including for and sampling uncertainties can be accounted for (Lynch et al. 2014; Kofler et al. 2011). Pooled sequencing has been successfully used to study population evolution in, for example, quail (Boitard et al. 2013), drosophila (Zhu et al. 2012) arabidopsis (Fracassetti et al. 2015) and the wild tomato species *S. chilense* (Böndel et al. 2015). Here we show proof of principle that pooled RENSeq can be used to identify R-genes of interest within a single population.

Our main aim is to identify R-genes that maintain polymorphisms within wild populations. We provide proof-of-principle in *S. pennellii*. Due to its limited genetic diversity, *S. pennellii* is particularly suited to test the statistical power of various population genetics methods on pooled data. We accurately identify a large set of NLR genes in the species and provide robust analysis to identify SNPs and calculate population genetics statistics. With this, we show that a small subset of R-genes maintains particular high diversity within *S. pennellii*.

## Methods

### NLR identification, analysis and probe design

To identify, high confidence NLR genes, we used the published *Solanum pennelli* sequence data and NLRParser as recommended by the authors (Steuernagel et al. 2015) We ran MAST (Bailey et al. 2009)(e=10>-6) using previously described NLR-associated motifs (Jupe et al. 2012). Matching sequences were extracted and submitted to NLRParser for annotation. The output was used to extract gene sequences and gff files with predicted protein annotations, to be used in follow-up analysis. A phylogenetic tree based on protein alignment was constructed using the extracted NBARC domains of the identified NLR. All domains were aligned with MUSCLE (Edgar 2004). Manual curation and removal of the biggest gaps was done in jalview (Waterhouse et al. 2009) before construction of the tree with PhyML (Guindon et al. 2010) (WAG model, BioNJ starting tree and NNI tree searching, 100 bootstraps).

Probes (S_File 2) were designed using Agilent’s SureSelect Software with the predicted NLR for *S. pennellii* and published NLR for *S. lycopersicum*, *S. tuberosum* and *A. thaliana*. We also included a set of 22 control genes used in previous evolutionary studies of potato or tomato (S_File 3). These included five resistance signaling associated genes (Pto, Fen, Rin4, Prf and Pfi) (Rose et al. 2011), three proteases (Rcr3, C14 and PIP1) and 14 metabolism related genes, the so-called reference genes in Böndel et al. (2015). We used BLAST and a second run of NLRParser to confirm that all targeted sequences were indeed putative NLR genes. Several probes gave false positive hits (targeting LRR-containing, but non NLR genes). Those probes were manually removed. In total 12,331 probes were selected to use with the SureSelect platform.

### Plants, DNA extraction and RENSeq

Ten *S. pennellii* plants (LA0716) were grown in our glasshouse under 16 hr light conditions and a minimum temperature of 18 °C. The seeds were obtained from Wageningen University CGN. DNA was extracted using a CTAB method. The DNA was quantified using Life Technologies’ Qubit and quality confirmed with Agilent Bioanalyzer 2100. DNA for 10 plants was pooled and NLR enrichment was performed according to Agilents SureSelect XT protocol with minor modifications: DNA was sheared on a Covaris S220 to 800 bp, size selection and cleaning was done using AMPure XP beads (Beckman Coulter) in two steps using 1.9:1 and 3.6:2 fragment DNA to beads ratio. The quality was assessed using a Bioanalyzer 2100 (Agilent). End repair, adenylation and adaptor ligation were preformed as described by Agilent. Pre-capture amplification was done using Q5 high fidelity PCR mixes. The amplified library was quality checked on a Bioanalyzer 2100. Hybridisation was performed as suggested for libraries <3 Mb. The library was indexed with 8bp index primers using Q5 PCR mix and quality was assessed using the Bioanalyzer 2100 and quantified using Qubit. Our library was pooled with seven other samples in equal DNA amounts and the resulting pool was quantified by qPCR using the NGSLibrary quantification kit for Illumina (Quanta biosciences) and diluted down to a final concentration of 20 nM. Illumina MiSeq was run twice on the same library following the manufacturers instructions for MiSeq v3. chemistry.

### Data Analysis

Our SNP detection methods are outlined in detail in Figure S1. FASTA files with sequencing data were quality controlled (QC) using trimmomatic (Bolger, Lohse, et al. 2014)(HEADCROP:3 SLIDINGWINDOW:4:30 TRAILING:30 MINLEN:40) and mapping was performed with trimmed reads using Stampy (Lunter & Goodson 2011) and BWA (Li & Durbin 2009)(default settings). Figure S2 shows the quality scores before and after trimming. Low quality mappings and duplicated reads were removed using Picard Tools (http://broadinstitute.github.io/picard/), before SNP calling. SNP calling was performed using Popoolation (Kofler et al. 2011), using the authors recommended settings, min-cov was varied from 3 to 9 (Figure S3A) and the expected allele count set to 20. We tried several sub-sampling methods. Figure S3B shows that sub-sampling in general appears to reduce the number of called SNPs and does not improve the quality. In addition, we used GATK Haplotypecaller and SelectVariants (McKenna et al. 2010). GATK allows for advanced filtering options, hence we used filters based on our Sanger sequenced data as outlined in S_File 5. For completeness we used two more popular SNP callers Varscan2 (Koboldt et al. 2012) and BCFTools (http://www.htslib.org/) using default settings for polyploid organisms.

The classic population genetics statistic π (Tajima 1983), was computed based on the estimated minor allele frequencies using SNPGenie (Nelson et al. 2015). The folded Site-Frequency Spectrum (SFS) estimations were done using several methods. Pool-HMM (Boitard et al. 2013) was run to calculate the allele frequency in our data (option-spectrum) directly from the alignment file. This data was fed back into Pool-HMM (option-estim) to estimate absolute allele frequency and summarised into folded spectrum. Secondly, a SFS was calculated from GATK output (generated using HaplotypeCaller with-ploidy 20), by parsing expected allele frequencies from the filtered output VCF, folding and summarising them. Lastly, we used filtered Popoolation outputs and deduced SFS from the observed allele frequencies. We computed the ratio of non-synonymous to synonymous diversity πN/πs using SNPGenie which uses an estimator based on the method of Nei and Gojobori (Nei & Gojobori 1986). Possible homologues for all the SNP containing genes were identified using BLAST against the curated swissprot database, to allow identification of homologues of evidence based NLR. Only NLR with >30% sequence identity and over 70% coverage with the original NLR were reported.

### Sanger sequencing

Primers were designed to anneal around at least one exonic region of the following genes: Sopen02g021920, Sopen12g030570, Sopen11g028610 and Sopen12g032710 (S File 6). Genes were amplified from DNA extracted from each of the individual plants used in our pool with Q5 polymerase (NEB), using the manufacturers recommendations. Amplified gene-fragments were purified (Qiaprep Qiagen) and sequenced directly, or ligated into the pENTR-TOPO2.1 vector (Life technologies) and transformed into *E. coli* TOP10 cells. Positive colonies were selected and plasmid DNA was extracted using Qiagen Qiaprep.

To identify all SNPs at each gene segment, we sequenced at least two plasmids per plant. We used CodonCode Aligner (CodonCode Inc) to check the sequence quality and align the plasmid sequenced with the reference genes. Up to 21 SNPs were manually annotated for each gene section.

### Visualisation

Visualisation of reads, annotations, motifs and SNPs was done using IGV (available from Broad Institute). Mapped reads were shown on the reference sequence and bedtools was used to generate custom tracks for the different NLR motifs, gene annotations and SNPs. Graphs were made in R (R Foundation for Statistical Computing, Vienna, Austria), using the package ggplot.

## Results

### *S. pennellii* contains 220 high confidence NLRs

The automated gene annotation for *Solanum pennellii* (Bolger, Scossa, et al. 2014) contains 486 proteins that contain domains associated with canonical NLRs. However, annotations are rather incomplete and describe only individual domains (214 NB-ARC, 259 Leucine Rich Repeats, 13 CC, TIR or other domains). As individual NB-ARC or LRR domains can also be part of other signaling proteins, like Receptor-like Protease (RLPs), careful re-annotation was required. We reannotated *S. pennellii* proteins and inferred whether they were putative complete or partial NLRs. We ran NLRParser against the predicted proteins for *S. pennellii* V2. This yielded 220 putative NLRs, of which 93 were complete (S File 1). We found 164 members of the CNL class, 39 of the TNL class, 16 lacking their N-terminus. As in previous RENSeq studies (Andolfo et al. 2014; Jupe et al. 2013), manual inspection showed that some putative NLRs might be wrongly annotated in the *S. pennelli* V2 genome. Some of our reads aligned well outside the annotated genes. As we were not yet able to accurately predict coding regions laying within these reads, which will be required for calculation of population genetics statistics, these reads were ignored and we focused only on those NLRs for which coding region data was available. To show that our dataset is likely to be a good representation of the NLRs to be found in *S. pennelli*, we constructed a phylogenetic tree based on the NB-ARC domain of the identified NLR. Figure 1 shows that our tree contains the main NLR classes that can be found in other tomato species and close homologues of known NLRs from unrelated species, similarly to those described for *S. lycopersicum* and *S. pimpenellifolium*.

**Figure 1.**
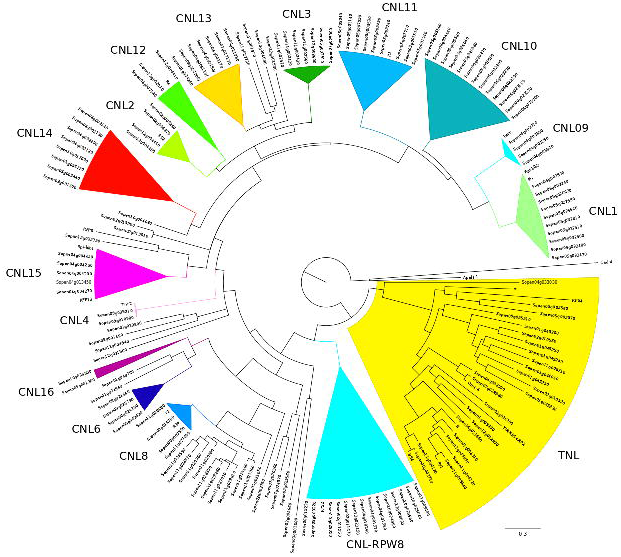
NLR genes in *S. pennellii*. Phylogenetic tree for the identified *Solanum pennellii* NLR genes generated using PhyML (WAG) with 100 bootstraps after alignment of all NB-ARC using MUSCLE. TNLs are highlighted in yellow background. Collapsed triangles represent known NLR clusters with high bootstrap values (>50). NLR families are indicated above the different clades and several named resistance genes from other species have been included for references.

### Sequencing, QC and mapping statistics

We used sequence data of the 220 predicted NLR together with previously annotated NLR from tomato (*S. Lycopersium*), potato (*S. tuberosum*) and previously described known NLR sequences (Jupe et al. 2012) to design NLR specific probes (S File2). DNA samples were sequenced as part of a larger pool. Two runs were done for our pool, which resulted in 805,122 and 2,147,039 reads. We performed basic quality control with Trimmomatic and trimmed all parts of the reads with quality lower than 30. Unpaired and low quality read pairs were removed and finally we retained 669,869 and 1,283,203 high quality paired reads. We were able to map 642,331 and 1,230,551 of the read pairs to the reference using Stampy for run1 and run2 respectively and 494,012 and 986,210 read pairs using BWA. Downstream analysis revealed that the BWA alignment, gave better results for the SNP calling, hence we thereafter report the values obtained with the BWA mapped reads only.

### RENSeq provides deep coverage in targeted regions

To assess the success of our enrichment sequencing, we plotted the depth of coverage per site against the fraction of the targeted region with the given coverage. Our probes were designed using exon data only, this reduces the coverage in intronic regions, but assures high read depth in coding regions. Figure 2 shows that close to 80% of the exonic target regions for the 22 control genes has a coverage of at least 130 reads, and 50% a coverage of at least 269 reads. For the NLRs, 80% of the predicted target region has a coverage of 245 or higher, and 50% a coverage of more than 408. As our initial mapping might contain misaligned or duplicated reads and mapping over introns, we performed an additional series of quality controls and filtering before identification of SNPs in both control and NLR datasets. Figure 1 shows the coverage plot after de-duplication and filtering. The coverage at the first quartiles (e.g. 75% of the regions with higher coverage) is 112, 172 and 251 x in respectively run1, run2 and both runs combined, whereas the median coverage was 163, 243 and 346 x respectively.

**Figure 2:**
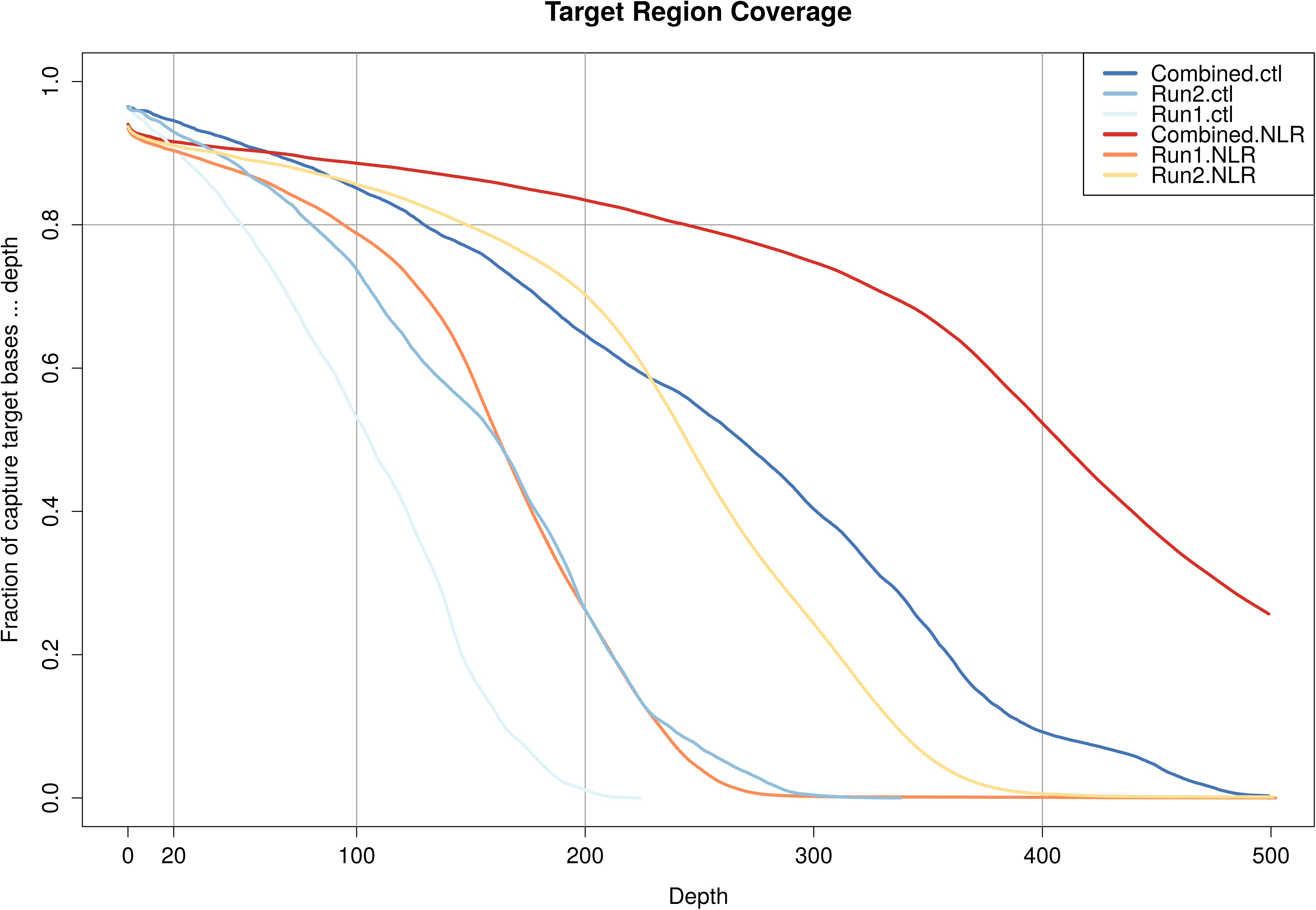
Coverage of targeted region. The fraction of bases in the targeted area having a coverage of a certain depth (x-axis) or deeper. The lines represent the individual runs and the combined data, separated for the NLR regions and the set of control (ctl) genes. The plot represents the data after preprocessing.

### GATK and Popoolation show highly congruent SNP calls in our population

Next we set out to identify SNPs in all exons of the NLR and control genes within our sequenced population. We ran Popoolation using different cut-off values to establish the maximum sensitivity while minimising the number of false positives. SNPs were called for run1, run2 and both runs combined, with minimum coverage set at 20, 30 and 40. Assuming equal amounts of DNA per plant and an average coverage near 120 in the run with the lowest coverage, we would expect a singleton allele frequency of 1/20. Minor singleton alleles should thus be readily picked up in the majority of cases with a minimum SNP count of five or six. Figure S3 shows that with low minor allele count (3-5) very large numbers of SNPs are detected, and that indeed after the count of six the detection curves flatten off. Importantly, differences between separate runs (and thus read depth) as well as the minimum overall depth tend to have a negligible effect on SNP calls (with mincount 5-9) (Figure S3). However, at higher stringency we observe a loss of sensitivity (mincount > 10). To guarantee high quality SNPs, we decided to keep the minimum depth for follow-up analysis at 30. This way, minor alleles occurring in frequency 4/20 can still be found with the minimum SNP count set at six. Lowering the minimum count could increase false positive rates in highly covered regions due to possible PCR bias. We also calculated the average coverage for all exons of each predicted gene to assure no correlation between SNP and coverage. Sub-sampling strategies implemented by Popoolation appear to have detrimental effect on the SNP calling (Figure S3B) and were not used. Using the setting described, in total 249 SNPs were identified in the NLRs.

Next we used GATK as a second method to verify the previously called SNPs by Popoolation. Using GATK we could predict 222 SNPs. We compared GATK predicted SNPs with our popoolation data. We found that 185 SNPs in 12 genes overlap between both datasets (Table 1). We manually inspected all SNPs called uniquely for GATK and found that 20 were called because they showed difference from the reference genome, but did not show polymorphism within the sample, three were called in low coverage (<30) regions and seven were called with by GATK with fewer than six occurrences of the SNPs. The final six are close to indel regions. To avoid false SNP calling, we excluded those regions in Popoolation. We also analysed all SNPs called only with Popoolation, and 28 appear to be on locations where also low quality reads can be found and four are near too high coverage regions (likely PCR bias). We could not observe any oddities for the other 32.

**Sheet 1.**
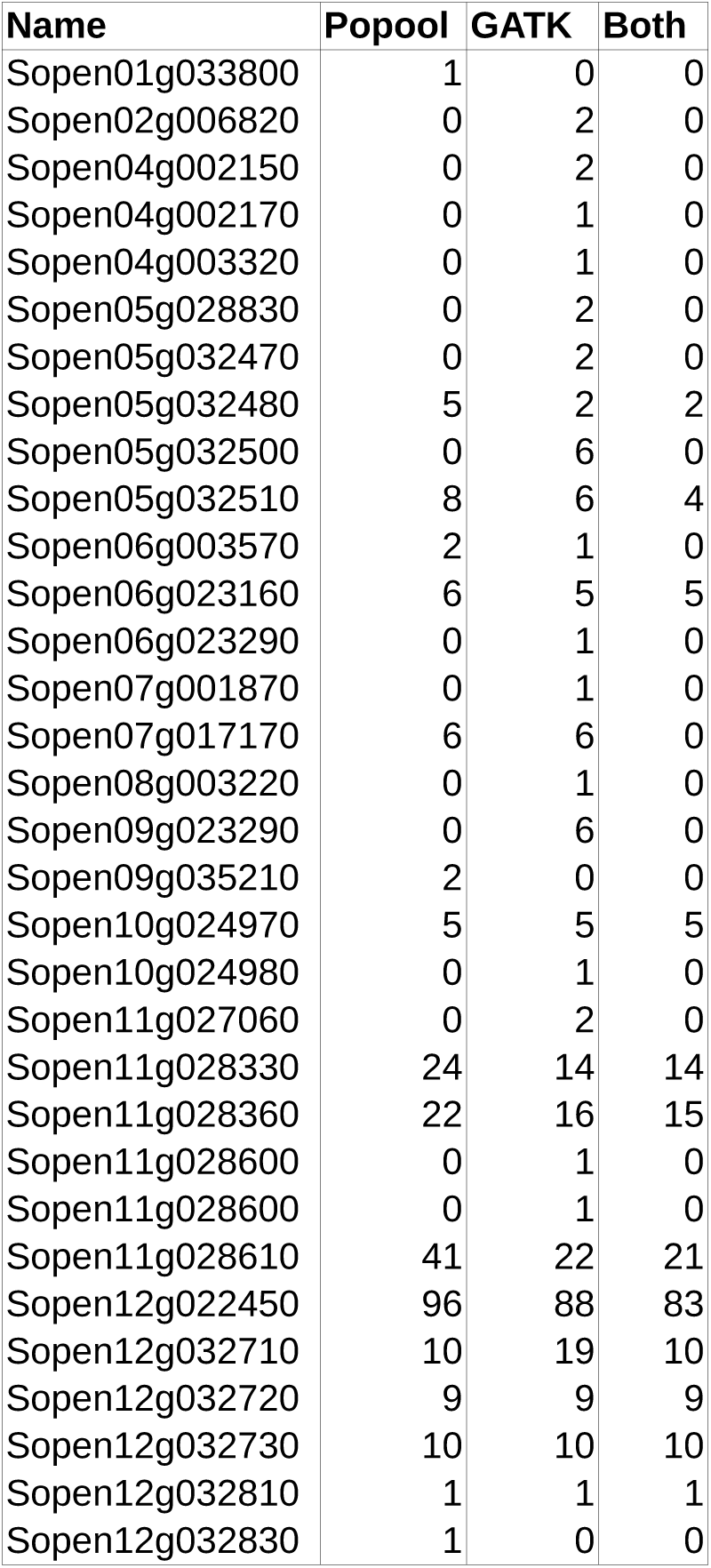

| Name | Popool | GATK | Both |
| --- | --- | --- | --- |
| Sopen01g033800 | 1 | 0 | 0 |
| Sopen02g006820 | 0 | 2 | 0 |
| Sopen04g002150 | 0 | 2 | 0 |
| Sopen04g002170 | 0 | 1 | 0 |
| Sopen04g003320 | 0 | 1 | 0 |
| Sopen05g028830 | 0 | 2 | 0 |
| Sopen05g032470 | 0 | 2 | 0 |
| Sopen05g032480 | 5 | 2 | 2 |
| Sopen05g032500 | 0 | 6 | 0 |
| Sopen05g032510 | 8 | 6 | 4 |
| Sopen06g003570 | 2 | 1 | 0 |
| Sopen06g023160 | 6 | 5 | 5 |
| Sopen06g023290 | 0 | 1 | 0 |
| Sopen07g001870 | 0 | 1 | 0 |
| Sopen07g017170 | 6 | 6 | 0 |
| Sopen08g003220 | 0 | 1 | 0 |
| Sopen09g023290 | 0 | 6 | 0 |
| Sopen09g035210 | 2 | 0 | 0 |
| Sopen10g024970 | 5 | 5 | 5 |
| Sopen10g024980 | 0 | 1 | 0 |
| Sopen11g027060 | 0 | 2 | 0 |
| Sopen11g028330 | 24 | 14 | 14 |
| Sopen11g028360 | 22 | 16 | 15 |
| Sopen11g028600 | 0 | 1 | 0 |
| Sopen11g028600 | 0 | 1 | 0 |
| Sopen11g028610 | 41 | 22 | 21 |
| Sopen12g022450 | 96 | 88 | 83 |
| Sopen12g032710 | 10 | 19 | 10 |
| Sopen12g032720 | 9 | 9 | 9 |
| Sopen12g032730 | 10 | 10 | 10 |
| Sopen12g032810 | 1 | 1 | 1 |
| Sopen12g032830 | 1 | 0 | 0 |

We further tested Varscan and Bcftools to call SNPs in our dataset, however both these callers seem to under-perform with 172 and 130 SNPs respectively. Possible reasons might be that contrary to Popoolation and GATK, the versions we used have not been optimised for multiploid (>2) specimens or pooled data. Figure 3A shows a Venn Diagram with the number SNPs called for each software. Popoolation and GATK together call the highest numbers of SNPs and also have the highest overlap.

**Figure 3:**
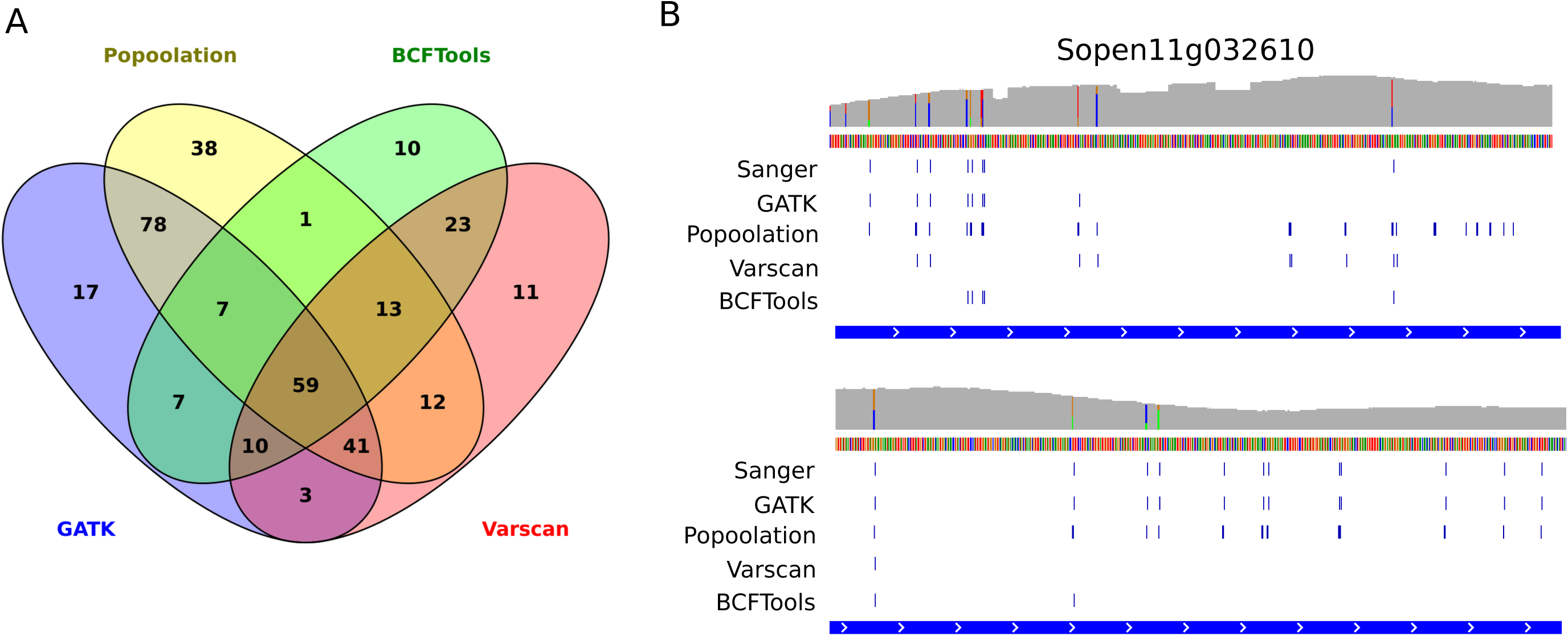
SNPs calls from four different callers. A) Overlap of called SNPs between different SNP-callers. Popoolation and GATK share the most common SNPs. B) SNPs called for a region of NLR Sopen11g028610. Top shows the coverage (grey) and SNPs that appear directly from the .bam file (including putative false positives). The blue lines in the lower parts of the figure show the SNPs as identified by Sanger sequencing and four SNP-callers. Popoolation and GATK show the best performance judging by overlap.

We also used Popoolation and GATK to identify polymorphisms in our control genes. Overall 12 SNPs were called in the control gene-set by both softwares, using settings previously described. One SNP was called by GATK only because it differed from the reference genome, but it did not show polymorphisms within our sample. Thus highlighting the importance of noting how SNP callers treat a reference sequence. As we are only interested in variation within our population (and not with the reference genome), such SNPs will be omitted in the remainder of this manuscript.

### All SNPs can be verified using Sanger sequencing

To verify our SNP calling using Sanger sequencing we designed primers annealing around one or more exons of two non-NLR genes, Sopen02g021920 (Rcr3) and Sopen12g030570 (C14) and two NLRs, Sopen11g028610 and Sopen12g032710 (S File 3). Our Sanger sequencing data confirm that Sopen02g021920 does not contain any polymorphisms (S File 6). For simple genomic regions, like those in Sopen12g030570 and Sopen12g032710, both GATK and Popoolation identified all Sanger sequenced SNPs. In complex regions, like part of Sopen11g028610, both GATK and Popoolation seem to call several, non-overlapping false positive SNPs (Figure 3B). Due to its more flexible filtering we are better able to approach the true SNP set using GATK, yet no filtering method keeps in all positives and filters out all false negatives. Again, Varscan and BCFTools, significantly under-perform in this gene. To assure high quality SNPs to calculate population genetics statistics, we will use SNPs as called by both GATK and Popoolation (Table 1). This overlapping set shows lower false positive (3,6%) and false negative rates (6,4%) compared to the Sanger data than the individual SNP sets and also removes SNPs picked up because they only differ from the reference (see previous paragraph).

### Low sequence diversity was already evident in the original population

Since we pick up low number of SNPs in our populations, we wanted to infer how the maintenance of the plants in various collections affected genomic diversity in the NLRs. *S. pennellii* is a facultative selfing plant, some loss of diversity can be expected. However, both the TGRC (UC Davis, USA) and CGN (Wageningen University, Netherlands) who maintained this population, confirm that since acquisition (by TGRC in 1958 and Wageningen from 1985) no more than 5-10 reproductive rounds have taken place and multiple plants were used in the process of multiplication. This reasoning is based on information provided by TGRC (R. Chetelat, pers. com.) and Wageningen University (W. v. Dooijeweert, pers. com.). We can therefore reconstruct the following population model. We assume an initial heterozygosity H_0_ which is defined here as the probability to sample two alleles which are different in a population (Charlesworth & Charlesworth 2010) at the time of sampling. If one plant was initially sampled, the first generation of multiplication by selfing decreases heterozygosity by half to a value of H_1_ =0.5H_0_. If two or more plants were sampled, and crossed to produce F1, a proportion 0.5s of heterozygosity is lost due to the selfing rate s, yielding H_1_ =(1-0.5s)H_0_. Subsequently, between eight and 12 diploid plants were produced every generation and crossed randomly in TGRC and CGN. In such randomly mixing population of size 2N=16 or 2N=24 chromosomes, the expectation for the decrease of heterozygosity between two consecutive generations (t and t+1) is H_t+1_=(1-1/2N)H_t_. At the time point of our sample, the number of NLR genes showing heterozygosity is H_sample_ = 13/220. Applying these formulae, we can estimate the initial heterozygosity after *t* rounds of mutliplication as H_0_=H_sample_/[H_1_(1-1/2N)^t^]. The initial proportion of heterozygote NLR loci in the initial wild population of *S. pennellii* would therefore be between H_0_=[0.17, 0.21] for s=1, and H_0_= [0.12, 0.14] for s=0.5, when assuming t=10 generations of multiplication. For convenience heterozygosity equates here with the proportion of polymorphic loci in our 220 NLRs with the population sample of 10 diploid plants (20 chromosomes). Increasing the number of initial plants, would lower the expected initial heterozygosity even more. Hence, we can conclude that *S. pennellii* LA0716 must have had very low original diversity with more than 75% of the NLR showing no polymorphims.

### Different site frequency spectrum estimators yield comparable results

We used different methods to estimate the site frequency spectrum of our NLR data. Pool-HMM (Boitard et al. 2013) calculates an allele frequency spectrum (SFS) directly from the mapped reads and uses this as a prior to estimate SNP frequency at a given location. We used GATK to infer allele frequency in HaplotypeCaller (using -ploidy 20), expected allele frequencies were then extracted after filtering. Lastly, we estimated allele frequency from the Popoolation output data on minor alleles in our dataset. All individual SNP frequencies were summed and turned into a folded SFS of the population. Figure 4A shows that in absolute values, Pool-HMM shows many more singletons and overall SNPs in the data, but this is likely due to the absence of the necessary filtering options. The relative SFS calculated from Pool-HMM and GATK derived data show very strong congruence (Pearson correlation = 0.98).

**Figure 4:**
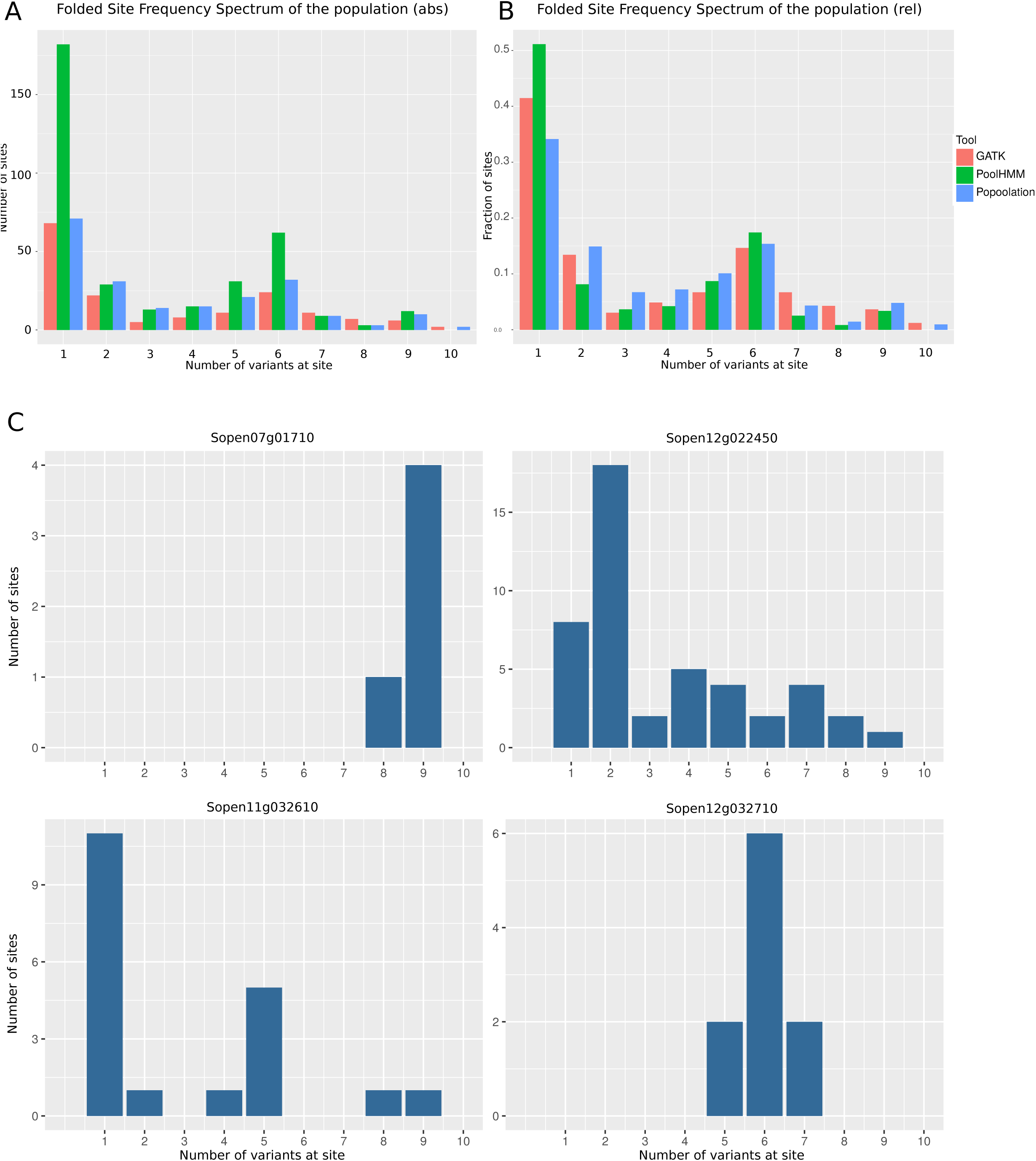
Site frequency spectra. Folded Site Frequency Spectra for the SNPs detected in our NLR set. X-axis shows number of variants per site, with 10 equals a frequency of 0.5 in our population. A) absolute folded SFS; y-axis shows actual number of sites. B) Relative folded SFS, y-axis shows the fraction of sites. C) Absolute folded SFS per gene.

Interestingly, our folded SFS shows an increase for class five to seven. Inspection of SFS per gene, reveals that, due to the low number of SNPs in our data, single genes with outlying SFS can be responsible for this pattern. Individual patterns for some R-genes show that indeed the genes seem to have differing spectra (Figure 4B). Sopen12g022450 shows an expected spectrum with high singleton count and flattening tail. Sopen07g01710 shows an increase in SNPs with intermediate frequency (greater than eight), whereas Sopen12g032710 shows an odd pattern with many SNPs occurring five to seven times, hence causing this intermediate frequency increase in the global SFS.

### NLR show differential evolutionary patterns

None of our 14 house-keeping control genes show any polymorphisms. For the pathogen related control genes, only one out of 8 (Sopen12g030570) had a significant number of SNPs within our population and a Ts/Tv ratio of 2.33. We identified 235 SNPs in our NLR data, with an average Ts/Tv ratio of 1.13. These SNPs were concentrated in only 13 NLRs. Strikingly the numbers of SNPs per gene range from 1 to 66 and are not correlated to gene length or average coverage depth (r=0,42 and 0.16). All genes meet the minimum coverage criteria in over 88 %. Nucleotide diversity is measured within the population as π per site and per gene (Table 2). Variation in π per gene ranges in two orders of magnitude between the different NLRs.

**Sheet 1.**
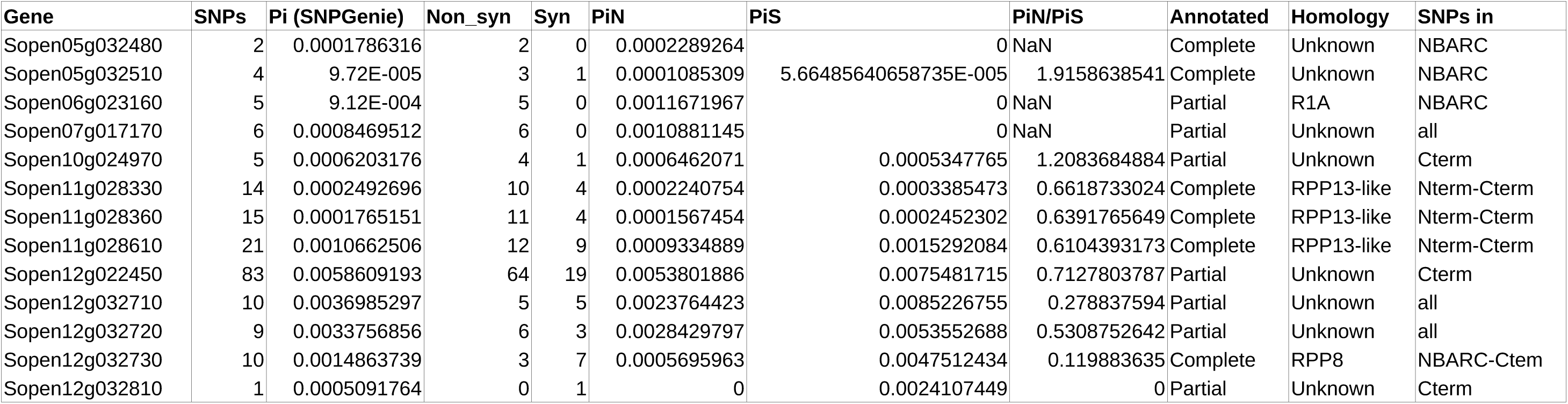

The assumptions that make Ka/Ks ratio a reliable estimator for selective pressure on R-genes between species, are not met when analysing data within populations (Kryazhimskiy & Plotkin 2008). To assess potential selective pressures we calculated π_*N*_/ π_*s*_ for all R-genes. (Table 2). In our set, overall, partial NLR genes show higher values for π_*N*_/ π_*s*_, however, many complete and partial NLR did not show any polymorphisms at all. Two NLRs (Sopen05g032510 and Sopen10g02490) show high (>1) π_*N*_/π_*s*_ values and three others (Sopen05g032480, Sopen06g023160, Sopen07g017170) contain several non-synonymous, but no synonymous mutations, both cases are indicative of positive selection.

Table 2 also shows that the identified SNPs are not limited to certain regions of the genes. Some NLR have SNPs in their C-terminus, other only in the NB-ARC domain or LRR domains, and in some cases SNPs are in all domains. Finally, we looked at the homology of our identified NLR with previously annotated NLRs from well known pathosystems. As expected with a highly divergent gene family, only five NLRs show resemblance with previously verified NLRs. These are one homologue of R1A from potato, one of A*rabidopsis* RPP8 and three of *Arabidopsis* RPP13.

## Discussion

We annotated NLR genes in a wild tomato species and show proof of principle that pooled MiSeq sequence data (250 bp reads) can be used to infer population genetics statistics to determine variation of R-genes within one small population of *S. pennellii*. Moreover, we show that even in populations with reduced diversity, large numbers of polymorphisms are maintained in certain R-genes.

### Identification of NLRs

We predicted 220 NLR genes in *S. pennelli*, which is an improvement over the previous annotation. This number is smaller than in cultivated tomato *S. lycopersicum* (326) and another wild relative *S. pimpenellifolium* (355) (Andolfo et al. 2014). Distribution amongst CNL and TNL classes is similar compared to both tomato species. Using current data, we find 93 NLR (43%) to be putatively full length genes. In cultivated tomato this number is about 70%.

Aforementioned studies on tomato, showed that so far only by manual curation and comparison with RENSeq sequence data one is able to identify all possible NLR-like regions on the genome. Unfortunately this comparison will not allow accurate annotation of open reading frames and we consider it outside the scope of this paper to perform and optimise such annotations. Moreover, increased NLR number in tomato and potato mainly result in additional partial genes and increases the number of complete NLRs by 17% only (Andolfo et al. 2014). Hence, our results indicate that in *S. pennellii* fewer NLRs are present. The phylogenetic reconstruction of the NLR family shows that our set of NLR genes covers the breath of NLR families observed in other *Solanum* spp. and we are confident that we have not missed any known NLR family. Therefore, the difference in NLR numbers could be caused by the habitat of *S. pennellii*, which is relatively arid and where one could assume a lower pathogen pressure than for exampel for S. *pimpenellifolium* (Caicedo & Schaal 2004).

### Successful deep sequencing using few resources

We showed that using RENSeq, we can cost and resource effectively, get a sufficient coverage over our target region using only 1/8th of an Illumina MiSeq lane. Köffler *et al*. (2012) suggested that for accurate pooled data processing very large numbers (>100) of individuals are needed to accurately capture all polymorphisms in the data set. They assume that in these cases on average each individual will be sequenced once or twice, with the high number of individuals making up for eventual bias due to sample preparation. This approach might be recommended for species where many individuals can be easily obtained like *Drosophila*, but is less feasible for larger species, or wild specimens, where collected samples might not contain that many individuals. We show that an alternative approach, using fewer samples, but assuring high coverage (on average >30 per diploid individual) can be as successful in identification of polymorphisms in a population. To assure the quality of the identified polymorphisms, we extensively tested four SNP calling packages and compared our data with selected genomic regions that were subjected to Sanger sequencing. The software Popoolation has been specifically designed for SNP calling in pooled samples of many individuals. We find that on our dataset Popoolation (Kofler et al. 2011) slightly overestimates the number of SNPs present in the data, possibly due to lack of filtering options to remove biases in read composition introduced as an artifact of library preparation. GATK (McKenna et al. 2010) allows for more stringent filtering, however no filtering thresholds could be identified so that GATK alone had the best result. This could be due to the nature of our data, which comes from enrichment sequencing and thus has very unequal coverage, big differences between introns end exons and hence various biases that we could not fully capture with the available filters. Two other SNP callers significantly underperformed on our data, possibly because these were not optimised for pooled or mutliploid samples. In the end, we obtained the best results by merging the results and accepting only those SNPs that were called by both GATK and Popoolation. This strongly reduced the number of false positive calls, but might mean that in some low coverage regions minor alleles will not be counted. Validation using Sanger sequencing on selected regions showed however that in those regions 93,6% of all SNPs have been positively identified and also that only 3,6% of the SNPs were not identified in cases where they should have been. Overall, this shows that by combining callers, we are able to get both high sensitivity as high accuracy.

### Identification of SNPs in samples with reduced diversity

Overall we identified very low numbers of SNP. This might be partly due to the stringency of the SNP calling, however Sanger resequencing of a number of genes did not yield any additional polymorphisms. The more likely explanation is the composition of the population. The sequenced plants come from a facultative selfing population collected in 1958 (Atico, Peru) and has been propagated during five to ten rounds at the TGRC and Wageningen University as small populations of eight to 12 plants (by pollen mixing and crossing). It is possible that the original population consisted of very few closely related specimens (maybe even one single plant) and that diversity has therefore been lost in the sampling and propagation processes. Our calculations show that the original proportion of genes with heterozygosity in the population could have been 10% or lower. With the current diversity found at 6% this shows that even though the multiplication and initial sampling have decreased heterozygosity in our NLR genes, the initial population exhibited very low genetic diversity to start with. This is consistent with the diversity of self compatible species to be much lower than that of self incompatible species. This is exemplified by the fact that using AFLP markers more diversity (75% polymorphic sites) was observed within one accession of self incompatible *S. peruvianum*, than between multiple accessions of self compatible *Solanum* spp. like *S. pimpenellifolium* (7%) (Miller & Tanksley 1990) Recent studies confirm such high levels of polymorphisms to occur only in self incompatible species (Städler et al. 2008).

We must note that the SFS will be strongly affected by genetic drift occurring during the multiplication process. This was seen in our global and per gene SFS with an excess of intermediate frequency variants. However, the genes we found to be polymorphic in our sample, will have been diverse in the initial population due to possible past selective events and provide an insight in the number and location of polymorphisms in different genes.

### Maintained polymorphism in C14 and NLR genes

We can identify polymorphisms in our control gene, C14. C14 is a tomato protease targeted by multiple effectors from *Phytophthora infestans*. It has been shown to be under diversifying selection in wild potato (Kaschani et al. 2010). This does not seem to be the case in several wild tomato species (Shabab et al. 2008), which are thought not to be a natural host for *P. infestans*. Also in our population, C14 polymorphisms are predominantly synonymous and we detect no sign of diversifying selection. Interestingly, we did not identify any SNPs in another protease, Rcr3, which is under balancing selection in *S. peruvianum* (Hörger et al. 2012). Also, Pto, Fen, Rin4, Prf and Pfi do not show polymorphism either, though they have been shown to be under selective pressure in S. peruvianum (Rose et al. 2007, 2011).

We identify after filtering 13 NLRs with one or more polymorphisms. Based on our above computations, we expect that heterozygosity at these genes reflects ancestral polymorphism in the initial population. These genes may thus show adaptation to different selective pressures, that could be caused by absence of or presence of certain pathogens on this specific population. Previous data from *Arabidopsis* suggests that when comparing different NLRs within a given genome, heterozygosity is larger in LRR regions (Mondragón-Palomino et al. 2002). However, we find no evidence that within one NLR polymorphisms between individuals are restricted to a certain region of the gene. This may be partly due to our current dataset containing too few SNPs in too few genes to identify trends and link selection pressures on the genes to the place or domains where the domains occur.

Five NLRs in our dataset show higher a higher π_*N*_ than π_*s*_ value, indicating possible positive selection. Due to the low diversity of our sampled population, we acknowledge that a high π_*N*_/π_*s*_ ratio however, does not as such suggest high positive selection pressure. As such, within gene diversity could be a better indicator for evolutionary pressure in this population, because this could be a sign of balancing selection. In terms of polymorphisms, certain individual genes indeed stand out. One of the genes that has maintained the highest number of polymorphisms within our population (Sopen11g028610), is an ortholog of *Arabidopsis* RPP13. RPP13 is known to maintain extreme high numbers of polymorphisms in wild populations (Rose et al. 2004), which is congruent with the highly polymorphic nature of its recognised effector Atr13 (Rentel et al. 2008; Sohn et al. 2007; Leonelli et al. 2011) and likely loss of fitness in the wild when one or multiple allelic variant disappear from the population. The highest number of polymorphisms can be found in Sopen12g022450. With 83 putative SNPs all in the Leucine Rich Repeats of the gene. It must be noted that this gene has been annotated as ‘partial’ gene and might not be functional. As with the previous example, it would be interesting to know Sopen12g022450 has a function in resistance and if the variants of it are maintained within different populations.

### Unraveling short-term NLR Evolution

A next step would be to test whether detected NLR variants show (partial) redundancies in terms of recognition. In grasses a number of resistance genes from fast evolving classes and classes with orthologues in 4 species have been cloned in rice and tested if they conferred resistance to 12 rice blast pathogen *Magnaporthe oryzae* strains. 15 out of 60 genes appear functional and no correlation was found between resistance and class or conservation between species (Yang et al. 2013) Resistances also appeared to be redundant between different pathogens, as observed in a larger study testing 132 NLR genes from cultivated rice. In the latter study 43% of the R-genes confer resistance against on average 2.4 of the 12 isolates tested (Zhang et al. 2015; Yang et al. 2013). Recent studies have shown how several NLR are required to work in pairs or networks, with closely related proteins sometimes conferring different functions (Eitas & Dangl 2010). Moreover, many NLR seem to be highly expressed also in susceptible interactions and NLRs can even be contributed to quantitative resistance effects (Corwin et al. 2016). Thus, analysis of long term evolutionary history using phylogeny would reveal only little about the recent selective pressures, state and activity of the NLRs.

As plants and pathogens are thought to adapt to one another within and between populations, our method can be used to identify NLRs that are under acute evolutionary pressure (see also Rose et al. 2007; and theory in Tellier et al. 2014). This is illustrated here as the identification of *S. pennellii* genes that maintained polymorphisms in our low-diversity population, including an RPP13 homolog. Follow-up work could include sequencing of multiple diverse populations, to help to identify functional R-genes from wild species, e.g NLRs under selective pressure. These methods can be expanded to polymorphism pathogen data, which will provide tests for current coevolutionary models (Tellier & Brown 2007; Tellier et al. 2014). To understand R-gene variation within and between populations of the same species, might help understand disease resistance ranges in crops and could solve questions on the molecular basis on non-host resistance (Stam et al. 2014). It will help define the durability of certain resistance genes and will hence be beneficial for future resistance breeding programmes.

## Acknowledgements

We thank Dr. Mareike Wenning and Christopher Huptas (TUM, ZIEL) for their assistance with MiSeq sequencing, Chase Nelson (U. South Carolina) for his support with the SNPGenie software and Hakyung Li, Ludvig Larsson and Saurabh Pophaly for help with the bioinformatics and/or sequencing. RS is funded through the Alexander von Humboldt Foundation.

---

Figure S1: Methods

Schematic overview of the methods used in this study. SNPs were called and curated after read mapping with Stampy (1). Filters were optimised based on Sanger data (2), when still large numbers of SNPs were not verified with Sanger sequence data, read mapping was repeated (3) with BWA, filtering was once more optimised (4).

Figure S2: Data Quality

Quality scores for sequencing data as reported by FastQC, before and after read trimming. Showing quality scores (y-axis) per read position (x-axis). Reads were trimmed to not get quality scores below 30.

Figure S3: SNP calling with Popoolation.

A) Total numbers of SNPs called (y-axis) against the stringency (minimum occurrences of a SNP before calling it). Low stringency (<5) gives large between run (e.g. coverage dependent) variation. B) Number of SNPs identified (y-axis) when subsampling to different depths (x-axis) for both runs combined (red) and each run individually (green, blue). Each graph represents a different sub-sampling method.

